# Transcriptional landscape of cell lines and their tissues of origin

**DOI:** 10.1101/082065

**Authors:** Camila M. Lopes-Ramos, Joseph N. Paulson, Cho-Yi Chen, Marieke L. Kuijjer, Maud Fagny, John Platig, Abhijeet R. Sonawane, Dawn L. DeMeo, John Quackenbush, Kimberly Glass

## Abstract

Cell lines are an indispensable tool in biomedical research and often used as surrogates for tissues. An important question is how well a cell line’s transcriptional and regulatory processes reflect those of its tissue of origin. We analyzed RNA-Seq data from GTEx for 127 paired Epstein-Barr virus transformed lymphoblastoid cell lines and whole blood samples; and 244 paired fibroblast cell lines and skin biopsies. A combination of gene expression and network analyses shows that while cell lines carry the expression signatures of their primary tissues, albeit at reduced levels, they also exhibit changes in their patterns of transcription factor regulation. Cell cycle genes are over-expressed in cell lines compared to primary tissue, and they have a reduction of repressive transcription factor targeting. Our results provide insight into the expression and regulatory alterations observed in cell lines and suggest that these changes should be considered when using cell lines as models.

**Highlights:**

- Cell lines differ from their source tissues in gene expression and regulation
- Distinct cell lines share altered patterns of cell cycle regulation
- Cell cycle genes are less strongly targeted by repressive TFs in cell lines
- Cell lines share expression with their source tissue, but at reduced levels

## Introduction

Cell lines are an essential tool in cellular and molecular biology, providing a lasting resource, ideally matching a particular genotype and phenotype in a controllable and reproducible setting. Cell lines have accelerated investigation of many biological processes, benefiting drug development and biomarker identification (Hu et al., 2006; Tan et al., 2011). Despite their merits as an experimental system, cell lines do not capture tissue complexity and heterogeneity, mainly because they consist of a single cell type that is adapted to grow in culture and lacks interactions with other cell types, the extracellular matrix, or paracrine signaling. There are many other factors that can influence cell line properties, including the methods that are used to establish and maintain them. Normal cells in culture divide a limited number of times and reach a state of nonproliferation called senescence; these are known as finite cell lines (Stacey, 2006). However, some cells acquire the ability to proliferate indefinitely in a process called transformation, which can be spontaneous or induced, and these are known as continuous cell lines.

Normal human fibroblasts are a common type of finite cell line that are easily isolated and grown in culture, but almost never spontaneously immortalize (Sherr and DePinho, 2000; Wright and Shay, 2000). Because normal fibroblasts show no genetic alterations in oncogenes and tumor suppressors, they are extensively used as model systems. For example, they have been used to better understand the mechanisms associated with the development of diabetic nephropathy, and to identify patients at risk of developing diabetic renal disease (Millioni et al., 2012).

Epstein-Barr virus (EBV) transformed lymphoblastoid cell lines (LCLs) are among the most widely created, archived, and analyzed continuous cell lines. LCLs have been extensively genotyped and sequenced as part of large collaborative projects, such as the International HapMap (International HapMap Consortium, 2003), 1000 Genomes (McVean et al., 2012), ENCODE (Dunham et al., 2012) and Genotype-Tissue Expression (GTEx) (Lonsdale et al., 2013) projects. EBV transforms resting B lymphocytes into proliferating lymphoblastoid cells, which carry multiple extrachromosomal copies of the viral episome and constitutively express six EBV nuclear antigens (EBNAs) and three latent membrane proteins (LMPs) (Young and Rickinson, 2004). Besides being used for genetic association studies, LCLs are also used to search for variants influencing gene expression (eQTLs), to measure response to radiation, and to test chemotherapeutic drugs (Jen and Cheung, 2003; Shukla et al., 2008; Watters et al., 2004). Despite their broad use, there has been concern about using LCLs as a model for primary tissues, with two small-scale studies finding differences in gene expression profiles between LCLs and primary B cells (Caliskan et al., 2011; Min et al., 2010).

While previous studies have analyzed the alterations in cell lines at the level of individual genes, analyzing cell line expression in the context of other sources of regulatory information may help us understand differences in gene regulatory processes. Complex cellular processes are defined by a combination of signaling pathways and cell type-specific regulators that may be represented in networks. By exploring regulatory networks, it has been possible to uncover patterns of transcriptional regulation specific to different cell types, such as immune (Heinz et al., 2010; Vandenbon et al., 2016) and stem cells (Müller et al., 2008), and to better understand cell differentiation (Yun and Wold, 1996), pluripotency (Kim et al., 2008), and development (Davidson et al., 2002). Moreover, commonly expressed transcription factors (TFs) may display distinct regulation patterns within different cell types (Neph et al., 2012) or different disease state (Glass et al., 2015).

The GTEx consortium generated a large multi-subject data set that offers an unprecedented opportunity to understand how well a cell line’s regulatory processes recapitulate those of its tissue of origin. GTEx version 6.0 includes RNA-Seq data for 127 paired LCL and whole blood samples, and 244 paired fibroblast cell line and skin samples.

A detailed investigation of gene expression and gene regulatory networks using these data indicates that, although many pathways are preserved between cell lines and their tissues of origin, some biological processes that help define the function of the primary tissue are altered. We also find that the cell lines exhibit changes in their apparent patterns of TF regulation. For example, while cell cycle genes are over-expressed in cell lines compared to their tissues of origin, they have an overall decrease in negative regulation by several TFs that are known to function as repressors. These findings suggest that the properties of cell lines must be carefully considered before using them as surrogates for primary tissue as some critical features of gene regulation may be missing or altered.

## Results

### Transcriptional landscape of cell lines and their tissues of origin

The GTEx version 6.0 project collected post mortem biopsies from multiple tissues and created LCLs and fibroblast cell lines. For the analysis described here, we used only data from research subjects for whom primary tissue and matching cell lines were available. Data were available for 127 paired whole blood samples and LCLs, and for 244 paired full-thickness skin biopsies and primary fibroblast cell lines (Lonsdale et al., 2013); 89 subjects have data across all four groups.

We found that cell lines and tissues have generally similar RNA-Seq transcriptomic profiles in terms of numbers of genes and the general functional categories to which those genes map (protein coding, antisense, pseudogene, lincRNA, and other; Figure 1A). We used principal components analysis (PCA) on paired samples between the two tissues and cell lines to examine the relationship between the expression levels of genes in cell lines and their tissues of origin. As seen in the projection of the first two components, the gene expression easily distinguishes the four groups (Figure 1B). The first principal component and the majority of the variability (37%) separated blood and LCLs from the skin and fibroblast samples. The second component (22%) separated tissues from cell lines. This indicates that while gene expression separates samples based on their tissues of origin, there is also a significant separation between cell lines and primary tissues.

**Figure 1.**
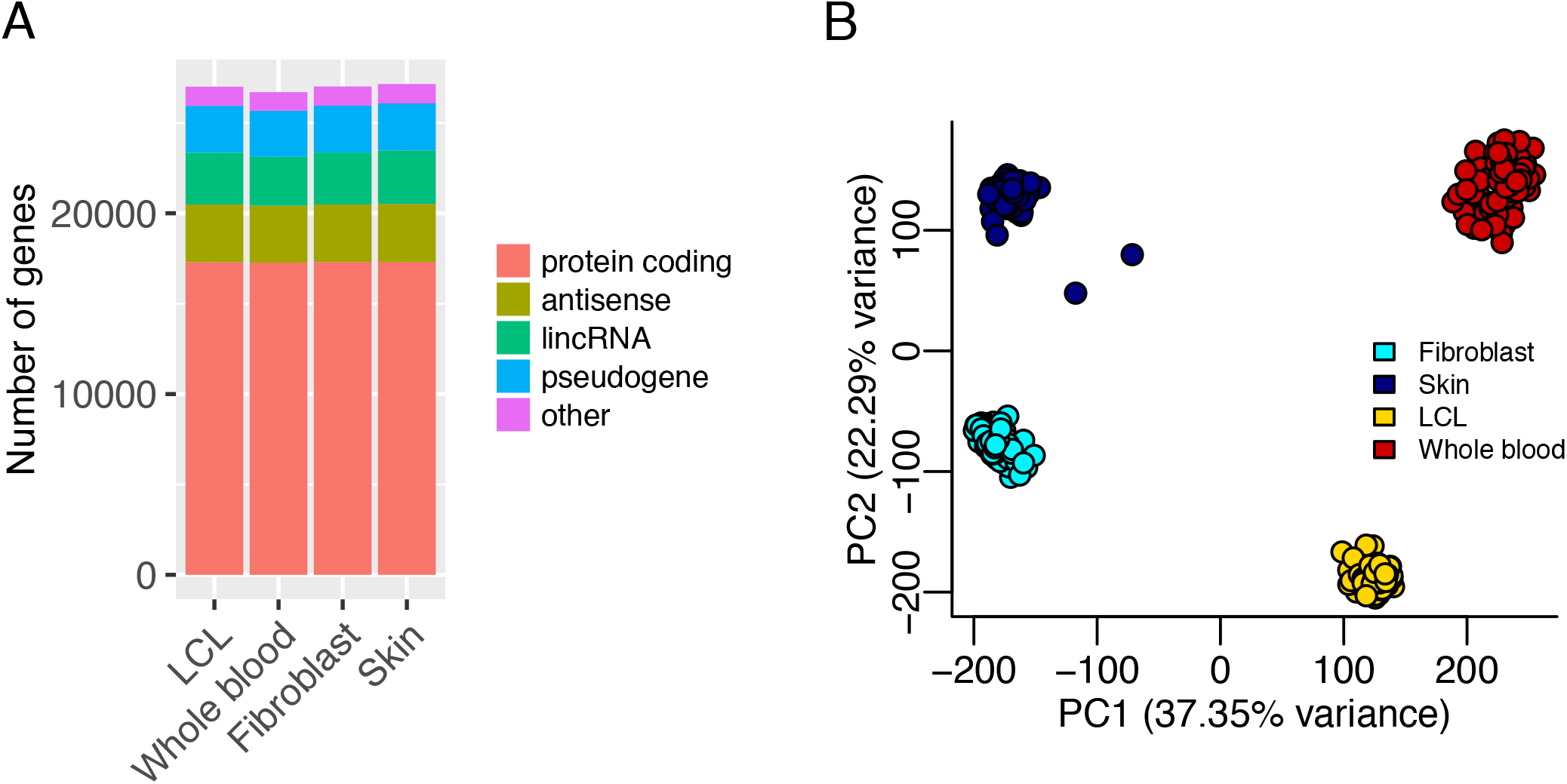
Similarity between cell lines and their tissues of origin based on gene expression. (A) Number of genes identified for each group (LCL, whole blood, fibroblast, skin). Genes were separated into biological classes using the definitions from GENCODE release 19 (GRCh37.p13). (B) Principal component analysis (PCA) of paired samples between the two tissues and cell lines (total of 89 subjects with all four samples) based on the normalized expression of all genes. The primary axis separates samples by tissue; the secondary axis separates primary tissue from cell lines.

### Differential expression between cell lines and their tissues of origin

To investigate which biological pathways are enriched in differentially expressed genes between cell lines and their tissues of origin, we used voom (Law et al., 2014) followed by Gene Set Enrichment Analysis (GSEA) (Subramanian et al., 2005). We found 8,617 genes (32%) to be differentially expressed between LCLs and blood paired samples (absolute log_2_ fold change > 2 and FDR < 0.05) with most of the differentially expressed genes (71%) over-expressed in LCLs (Figure 2A, Table S1). In comparing fibroblasts and skin paired samples, we identified 5,655 differentially expressed genes (21%). In contrast to the LCL-vs-blood comparison, most of the differentially expressed genes (68%) had increased expression in the primary tissue rather than in the cell line. Previous studies have also shown that approximately 30% to 50% of genes have significant differential expression between cell lines and tissue samples (Caliskan et al., 2011; Sandberg and Ernberg, 2005).

**Figure 2.**
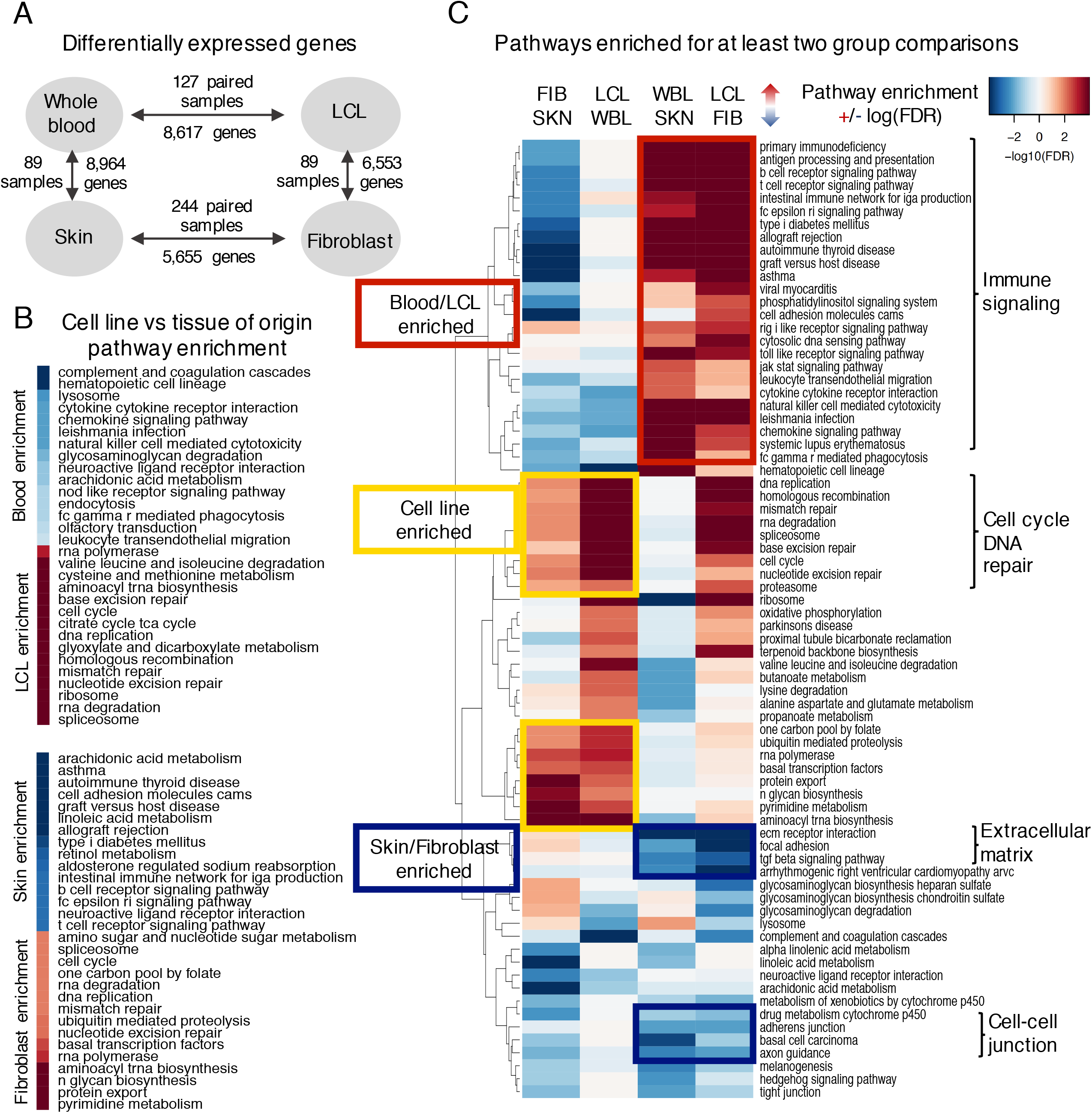
Pathways are differentially expressed between cell lines and their tissues of origin. (A) Number of differentially expressed genes (absolute log_2_ fold change > 2 and FDR < 0.05) using voom on paired samples. (B) Results of GSEA reported based on the log_10_(FDR) significance scale, with one group in red and the other one in blue. The 15 pathways most significantly different between one cell line and its tissue of origin. (C) Pathways enriched for at least two group comparisons (FDR < 0.05). The pathways differentially expressed between the tissues that are also differentially expressed between the cell lines (preserved pathways) are highlighted in red and blue. Pathways over-expressed in both cell lines compared to their tissues of origin are highlighted in yellow. See also Tables S1 and S2.

Using GSEA, with genes ranked by the moderated *t*-statistic from voom, we identified KEGG pathways (Kanehisa et al., 2015) enriched for differentially expressed genes between cell lines and their tissues of origin. Consistent with the separation observed in the PCA, both cell lines exhibit enrichment for pathways with similar biological functions compared to their tissues of origin (Figure 2B). While immune processes are down-regulated in cell lines, the pathways with positive enrichment are generally associated with cellular growth, and include cell cycle, DNA replication and repair, and transcription processes. Cell cycle genes have been previously found to be over-expressed in many cell line models (Lukk et al., 2010; Pan et al., 2009; Sandberg and Ernberg, 2005).

When comparing blood to LCLs, we found that pathways enriched in blood were related to immune system function, including complement and coagulation cascades, hematopoietic cell lineage, chemokine signaling, and natural killer cell mediated cytotoxicity, while the pathways enriched in LCLs were associated with cell growth and death, DNA replication and repair, transcription, translation, amino acid and carbohydrate metabolism (FDR < 0.05, Figure 2B). Similarly, when comparing skin to fibroblasts, the pathways enriched in skin were related to the immune system, lipid and vitamin metabolism, cell adhesion, and melanogenesis, while the pathways enriched in fibroblasts were associated with cell growth and death, DNA replication and repair, transcription, and protein degradation (FDR < 0.05, Figure 2B). The significance of all KEGG pathways is listed in Supplemental Table S2.

We also performed differential expression and KEGG pathway enrichment analysis comparing the two tissues and comparing the two cell lines. We found a number of immune signaling pathways enriched in blood compared to skin and in LCLs compared to fibroblasts (Figure 2C). For example, pathways related to the biological function of B cells (B cell receptor signaling, toll-like receptor signaling, antigen processing and presentation) were enriched in LCLs and blood when comparing them to fibroblasts and skin, respectively. However, some immune related pathways, including chemokine signaling and natural killer cell mediated cytotoxicity, were also enriched in LCLs and blood compared to fibroblasts and skin, but they were expressed at lower levels in LCLs compared to blood. The pathways enriched in fibroblasts and skin compared to LCLs and blood were associated with biological processes related to maintaining skin structure and organization and included cell-cell junction, extracellular matrix interaction, and TGF-β signaling (Figure 2C).

Overall, we found that the preserved pathways in cell lines are mainly related to the cell type specific functions (B cells or fibroblasts) rather than tissue-enriched functions. Further, many of the genes in pathways that help define the function of the tissue are under-expressed in cell lines relative to their tissues of origin.

### Cell line and tissue-specific gene regulatory networks

Understanding the structure of gene regulation in cell lines compared to their tissues of origin has the potential to help interpret differential expression results and to reveal important regulatory differences. We used PANDA (Glass et al., 2013) to integrate gene expression, predicted TF binding location and TF-TF interactions, and to estimate gene regulatory networks in LCL, blood, fibroblast, and skin (Figure S1A).

In the PANDA gene regulatory networks, an edge connects a TF to a target gene, and the associated edge weight indicates the strength of the inferred regulatory relationship. For each TF we computed the difference between the “out-degree” (sum of edge weights from that TF) in the cell line network and the corresponding tissue of origin network. We ranked TFs by their absolute difference in out-degree (differential targeting) and found that TFs with the largest differential targeting were involved in cellular responses to stress and DNA damage, and in the control of cellular growth (Figure S1B, Table S3). For both the LCL-vs-blood and fibroblast-vs-skin comparisons, many of the top differentially-targeting TFs, such as TP63, TOPORS, and KLF15, belong to the p53 family or interact with p53 and are important mediators of DNA damage response regulating cell cycle arrest, DNA repair and apoptosis (Candi et al., 2014; Haldar et al., 2010; Levrero et al., 2000; Lin et al., 2005).

We found SP1 and SP3 had increased targeting in cell lines in both network comparisons, LCL-vs-blood and fibroblast-vs-skin. These TFs have more than 12,000 binding sites in the human genome and are involved in essential cellular processes, including proliferation, differentiation, and DNA damage response (Beishline and Azizkhan-Clifford, 2015; Cawley et al., 2004). It is important to note that SP1 and SP3 are not differentially expressed between cell line and tissue samples. However, network comparisons captured the regulatory “rewiring” of these TFs and their target genes, revealing potential differences in targeting even in cases where the TFs themselves were not differentially expressed. Thus, these network models suggest that TFs alter their patterns of regulation in cell lines, either through changing their expression, altering the genes they target, or by participating in different regulatory contexts in controlling their potential target genes (Figure S1C).

### Cell cycle pathway genes are less strongly targeted by TFs in cell lines

We then tested whether or not changes in inferred TF targeting preferentially affected genes belonging to specific biological pathways. Similar to the TF’s out-degree, we calculated each gene’s “in-degree” as the sum of edge weights connected to a gene, which allows for a comparison of how strongly targeted each gene is by the complete set of TFs. We compared the in-degree differences for genes of a specific pathway against all other genes using an unpaired *t*-test (Table S4). For the pathways over-expressed in the cell lines, such as cell cycle, DNA repair, and DNA replication, we found a marked reduction of targeting in cell lines compared to their tissues of origin (Figure S2).

To better understand these differences, we explored the network around 121 genes of the KEGG cell cycle pathway (Figure 3, cell cycle gene names listed in Table S5). When comparing the log_2_ fold change of the expression levels of these genes with their differential targeting, we found a negative correlation (−0.48 to −0.62) for many TFs, including SMAD5 (Figure 4A), E2F8, ZBTB14, ETV5, HLTF, USF1, IKZF1, and USF2 (Figure S3). This indicates that, even though the cell cycle genes are over-expressed, they are less strongly targeted by these TFs in LCLs compared to blood. To confirm this, we used a permutation analysis using random gene sets equal in size to the cell cycle gene set and all negative correlations were significant (FDR < 0.05).

**Figure 3.**
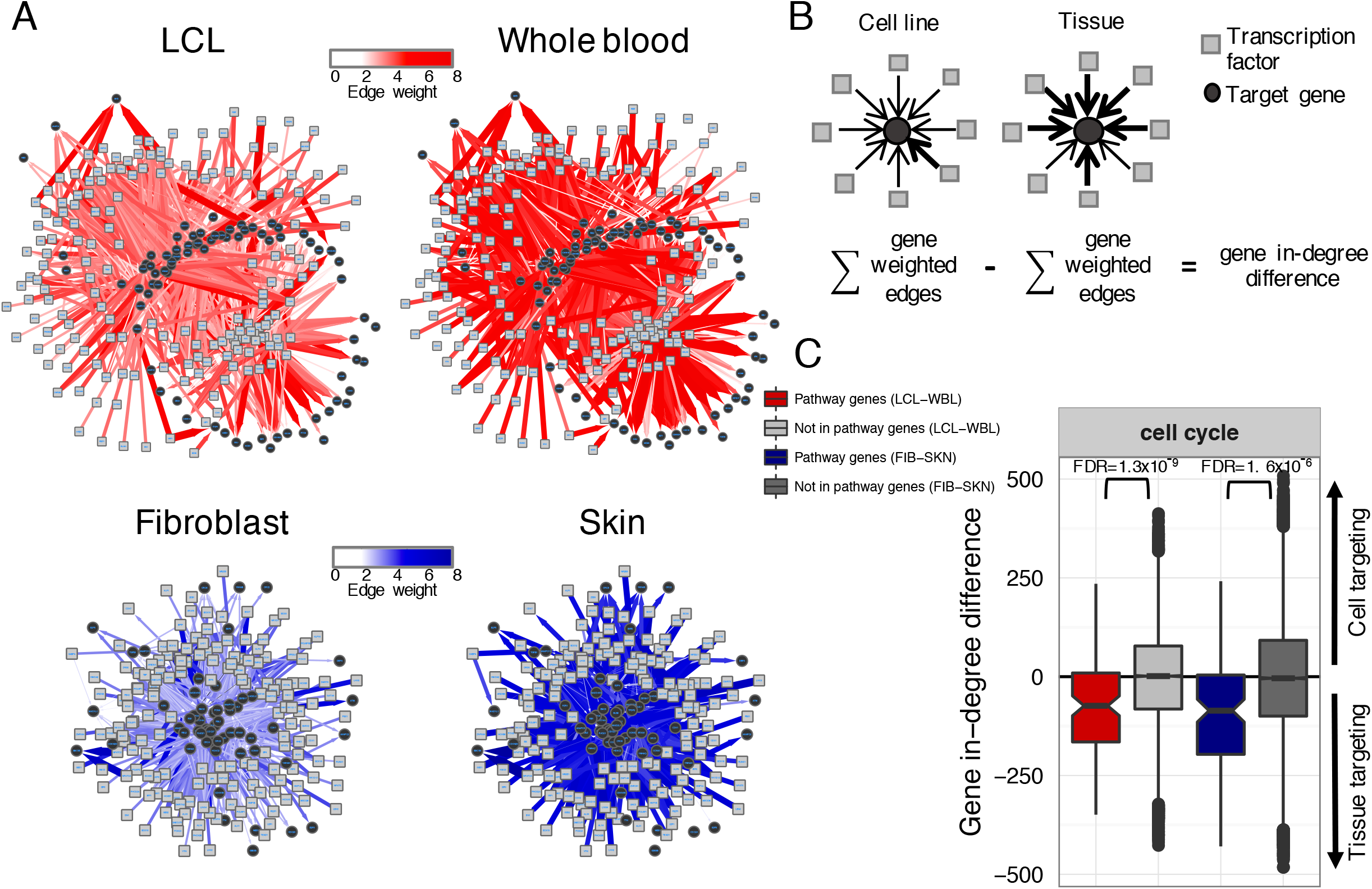
Cell cycle pathway genes are less strongly targeted by TFs in cell lines. (A) Group-specific gene regulatory networks were generated using PANDA. The illustrations represent subnetworks of the 1000 edges with the highest edge weight difference between a cell line and its tissue of origin around the cell cycle genes. The thickness and color indicate the edge weight strength between the TF and target gene (the edges shown have a weight greater than 2 in at least one network). (B) Illustration of the gene in-degree difference between each cell line and its tissue of origin. Positive values indicate higher targeting in cell lines, and negative values indicate higher targeting in tissues. (C) Boxplot of the gene in-degree differences for the genes in the KEGG cell cycle pathway and for genes not in this pathway (*t*-test). Reduction of gene in-degree difference indicates that the genes in the cell cycle pathway are less strongly targeted by TFs in the cell line compared to its tissue of origin. See also Figures S1 and S2, Tables S3 and S4.

This analysis suggests that these TFs play a role as negative regulators of the cell cycle. Indeed, these TFs are known regulators of the cell cycle, and many have documented roles in repressing genes that promote the cell cycle. Given the top TFs found in the network analysis as examples, SMAD5 can repress transcription, leading to proliferation inhibition after TGF-β signaling (Bakkebø et al., 2010). E2F8 directly binds to E2F family target genes and repress their transcription (Christensen et al., 2005; Ghazaryan et al., 2014; Li et al., 2008). ZBTB14 is a transcriptional repressor of the mouse *c-myc* gene (Numoto et al., 1993).

To corroborate our network predictions, we looked for other biological evidence that these TFs regulate cell cycle genes. We downloaded the LCL GM12878 ENCODE ChIP-Seq assays for the available TFs (SMAD5, IKZF1, USF1, USF2). We then identified the genes with peaks for these TFs in their promoter region. We also calculated the correlation between the expression of each TF and the cell cycle genes with TF ChIP-Seq binding evidence.

SMAD5, the TF with the highest inverse correlation between the expression and targeting of cell cycle genes (Figure 4A), binds to the promoter region of 113 out of the 121 cell cycle genes based on ENCODE ChIP-Seq data. As expected, we found a higher negative correlation between the expression of SMAD5 and the expression of its target genes in cell lines compared to tissue samples in GTEx (p-value = 8.7×10^−09^, Figure 4B). The combined GTEx/ENCODE results suggest that cell cycle regulation involves a complex interplay between changes in the expression of regulatory TFs and alterations in the binding of these TFs to their targets.

**Figure 4.**
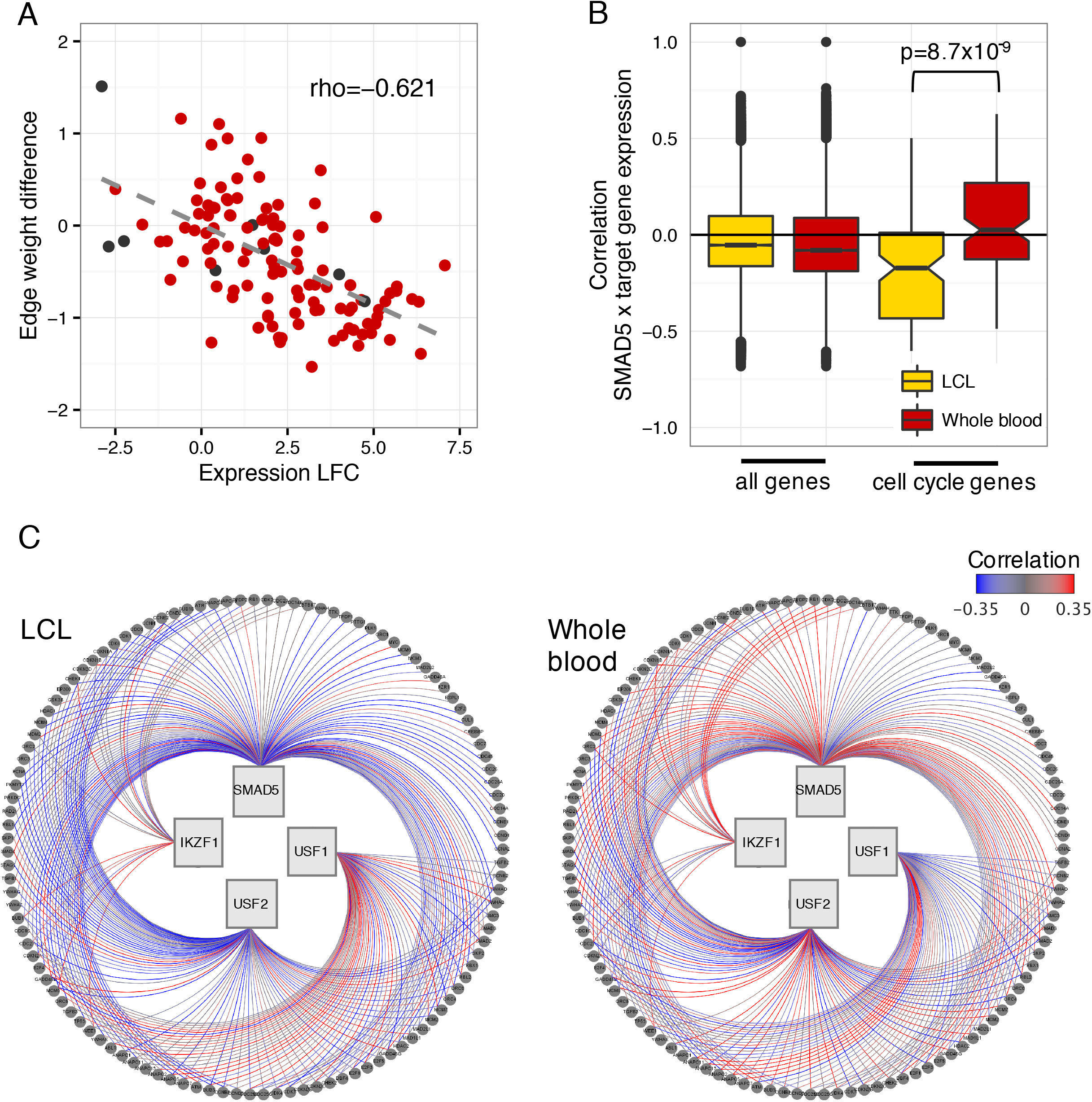
SMAD5 is differentially regulating cell cycle pathway genes. (A) Spearman correlation between the log_2_ fold change in gene expression (LCL-blood difference) of KEGG cell cycle pathway genes and the differential targeting they receive by the TF SMAD5. Red: evidence of SMAD5 ChIP-Seq binding on the promoter of the gene, black: no evidence of SMAD5 binding. The negative correlation observed indicates the cell cycle genes are more highly expressed but less targeted by SMAD5 in LCL compared to blood. (B) Boxplot of Spearman correlation coefficients between SMAD5 expression levels and expression levels of all genes, and between SMAD5 expression levels and the expression levels of cell cycle genes with SMAD5 ChIP-Seq binding evidence for LCL and blood samples. Difference in magnitude was tested using a Wilcoxon rank-sum test comparing LCL and blood. (C) Visualization of the correlation between TF and cell cycle gene expression for interactions that have ChIP-Seq binding evidence. More positively-correlated associations are shown in red, more negatively correlated are blue, and correlations near zero are gray. See also Figures S3, S4 and Table 5.

There is also extensive functional evidence that SMAD5 targets genes to inhibit cellular growth. SMAD proteins have a key role as signal transducers of the TGF-β family members to mediate growth inhibition and apoptosis (Derynck and Zhang, 2003). SMAD5 negatively regulates cell proliferation during embryonic hematopoiesis (Liu et al., 2003), in B-cell lymphoma (Bakkebø et al., 2010), and it induces cell cycle arrest in response to shear stress in tumor cell lines (Chang et al., 2008).

We repeated this analysis for the three other TFs with ChIP-Seq data available in ENCODE. We found similar trends for IKZF1 and USF2, but not for USF1 (Figure S3). Figure 4C shows a summary visualization of the expression correlation between the four TFs and the cell cycle genes with TF ChIP-Seq binding evidence, and supports the network-based conclusion that these TFs are negative regulators of cell cycle genes. The smaller correlation observed for USF1 might be due to the alteration of USF1 gene regulatory properties depending on its post-translational modification (Corre et al., 2009).

Experimental evidence also demonstrates anti-proliferative roles for USF1 and USF2. Loss or impairment of USF transcriptional activity is a common event in cancer cell lines and is associated with increased proliferation (Ismail et al., 1999; Qyang et al., 1999). Additionally, over-expression of USF, and in particular USF2, is known to suppress growth in a number of cell lines (Luo and Sawadogo, 1996; Olave et al., 2010). CDK4, which controls the progression of cells through G1, is transcriptionally regulated by USF1, USF2 and c-Myc in non-tumorigenic mammary cells (Pawar et al., 2004). However, in breast cancer cell lines USF is transcriptionally inactive, and the CDK4 promoter loses control by USF and c-Myc, altering CDK4 gene regulation and its ability to respond to signals in cancer cells (Pawar et al., 2004). We found similar differences in CDK4 regulation when we compared networks from LCL and blood. Consistent with its higher expression in LCLs, CDK4 is also less strongly targeted by USF1, USF2, and c-Myc.

When comparing fibroblasts and skin, we did not find the same strong negative correlation between cell cycle gene expression and specific TF targeting (Figure S4A). This may be due to the smaller changes we observed in expression of cell cycle genes in fibroblasts-vs-skin, in contrast to the LCL-vs-blood comparison, which is potentially related to the fact that LCL is transformed and fibroblast is not transformed. However, when we analyzed the relationship between SMAD5 and the cell cycle genes expression, we found similar results for fibroblasts (Figure S4B), indicating that some of the patterns we observe for LCLs may be true at a smaller scale in other cell line models.

## Discussion

Cell lines are widely used as experimental models to explore basic cellular biology, to study gene regulation, and to test drugs and other compounds. One important question is whether cell lines reflect the properties of the primary tissues from which they are derived. By studying gene expression and gene regulatory networks, we were able to uncover patterns of transcriptional regulation that differentiate cell lines from their tissues of origin. To the best of our knowledge, this is the first study that compares the differences in regulatory networks between cell lines and their tissues of origin and reveal differences not observed in differential expression analyses.

In comparing two tissue/cell line pairs, blood/LCLs and skin/fibroblasts, we find that while cell lines and their tissues of origin express similar numbers of genes and the functional categories to which those genes map are similar, there are also important transcriptional differences. We found that approximately 26% of the expressed genes were differentially expressed between the cell lines and the corresponding tissue of origin. Both LCL and fibroblast cell lines have increased expression of genes in pathways associated with proliferation and DNA repair relative to their tissues of origin. Cell lines also have reduced expression of genes associated with many of the primary functions that characterize the tissues. Our analysis shows that while many tissue-enriched pathways are downregulated in cell lines, cell type-specific pathways are preserved. This suggests that studies focusing on these cell type-specific pathways can use cell lines as models, although users should be aware of the unique aspects of cell line expression and regulation.

We investigated the potential drivers of the transcriptional changes by modeling gene regulatory networks and comparing the regulatory networks of cell lines and their tissues of origin. The most striking difference we observed was in regulation of processes associated with cellular proliferation. Processes including cell cycle, DNA repair, and DNA replication were more highly expressed in cell lines, and we found that they had lower overall targeting by a number of cell cycle-associated TFs that are known to function as repressors. These results indicate that cell lines switch off a number of transcriptional repressors that result in an overall increase in cell cycle-related transcription.

Consistent with our findings that LCLs lose overall negative transcriptional regulation, a recent study showed hypo-methylation of 250 genes after EBV transformation; in this case the cellular machinery could not maintain DNA methylation (Hernando et al., 2013). While alterations in the epigenetic profiles of LCLs have been demonstrated (Caliskan et al., 2011; Hernando et al., 2014), our analysis is the first to explore the changes of TFs regulatory targeting.

The fact that the GTEx cell lines were created in very different ways – one transformed and the other a primary cell line – suggests that the global alterations in transcriptional patterns may be associated with growing in culture, the lack of tissue context, and decreased cellular heterogeneity. Some of the changes may also be associated with the transformation process as we observed smaller changes in expression and regulation of cell cycle genes in fibroblasts compared to LCLs.

Understanding that differences exist between cell lines and tissues in patterns of TF targeting is important for designing and interpreting experimental studies using cell line models. This is especially true for the development of targeted therapeutics, where failures might be associated with therapeutics that target pathways that are altered in cell lines relative to the tissue from which they are derived. Our analysis provides insights into which transcriptional processes are altered and demonstrates that cell lines exhibit gene expression and regulatory changes that distinguish them from their primary tissues and that these changes and the processes associated with them should be considered when using cell lines as model.

## Author contributions

All authors contributed to the conception and design of the study. CLR and JNP analyzed the data. All authors contributed to writing and editing of the manuscript. All authors read and approved the final manuscript.

## Acknowledgments

This work was supported by grants from the US National institutes of Health, including grants from the National Heart, Lung, and Blood Institute (5P01HL105339, 5R01HL111759, 5P01HL114501, K25HL133599), the National Cancer Institute (5P50CA127003, 1R35CA197449, 1U01CA190234, 5P30CA006516), and the National Institute of Allergy and Infectious Disease (5R01AI099204). Additional funding was provided through a grant from the NVIDIA foundation. CML was supported by Sao Paulo Research Foundation (FAPESP) grant 2014/19062-9. This work was conducted under dBGaP approved protocol #9112.

## Experimental Procedures

### GTEx data

The GTEx version 6.0 RNA-Seq data set (phs000424.v6.p1, 2015-10-05 released) was downloaded from dbGaP (approved protocol #9112). Using YARN R package (version 1.0.0) we performed quality control, gene filtering, and normalization preprocessing (Paulson et al.). We identified and removed GTEX-11ILO due to potential sex misannotation. We grouped related body regions using gene expression similarity. For example, skin samples from the lower leg (sun exposed) and from the suprapubic region (sun unexposed) were grouped as “skin.” We filtered and normalized the data in a tissue-aware manner using smooth quantile normalization [github.com/stephaniehicks/qsmooth]. The final data set contains 549 research subjects (188 females and 361 males) comprising 38 tissues (which included two cell lines), 30,333 genes, and 9,435 samples. We filtered sex-chromosome and mitochondrial genes (retaining 29,242 genes).

We reduced the data set to only cell line and tissue-specific paired samples, which comprised 127 subjects with whole blood and EBV transformed lymphoblastoid cell line (LCL) samples, and 244 subjects with skin and primary fibroblast cell line samples; 89 subjects have data across all four groups. For the skin samples, an equivalent number of samples were obtained from the lower leg (n=123), and from the suprapubic region (n=121). We kept only the 27,175 genes with at least one TF binding motif in its promoter region (see section: Gene regulatory networks), so that we could use the same set of genes for differential expression and gene regulatory network analysis.

GTEx RNA-Seq version 6 was annotated using the GENCODE release 19 (GRCh37.p13). Thus, we defined the different types of genes (protein coding, antisense, pseudogene, lincRNA, and other) according to the same genome annotation downloaded from http://www.gencodegenes.org/releases/19.html.

### Principal Components Analysis

We performed principal component analysis (PCA) as implemented in the plotOrd function on the R package metagenomeSeq 1.12.1. PCA was applied to the full expression data matrix.

### Differential expression analysis

Differential expression analysis was performed using voom R package (version 3.26.9) (Law et al., 2014) and only paired samples between the two groups under comparison (LCL-vs-Blood, Fibroblast-vs-Skin, LCL-vs-Fibroblast, Blood-vs-Skin). Genes with Benjamini-Hochberg adjusted p-values of less than 0.05, and with absolute log_2_ fold change greater than 2 were considered to be differentially expressed.

### Pathway enrichment analysis

We performed Gene Set Enrichment Analysis (GSEA) to determine the biological functions related to the differential expression between cell lines and tissues (Subramanian et al., 2005). All genes were ranked by the moderated *t*-statistic produced by voom differential expression analysis. We used pre-ranked GSEA program (Java command line version 2-2.0.13) to calculate a running-sum statistic. We used the gene sets obtained from the KEGG pathway database that was downloaded from the Molecular Signatures Database (MSigDB) (http://www.broadinstitute.org/gsea/msigdb/collections.jsp) (“c2.cp.kegg.v5.0.symbols.gmt”). We performed 1000 gene set permutations to assess the statistical significance, and considered gene sets with FDR < 0.05 significant. We only considered gene sets of size greater than 15 and less than 500 genes after filtering out those genes not in the expression dataset, or 176 gene sets in total.

### Gene regulatory networks

We reconstructed gene regulatory networks using PANDA, a message-passing model that integrates multiple types of genomic data and infers the network of interactions between TFs and their target genes (Glass et al., 2013). PANDA starts with a prior regulatory network inferred by mapping TF binding sites to the genome, integrates protein-protein interaction and gene expression data to iteratively refine the network structure and deduces a final consensus regulatory network. In the regulatory networks estimated by PANDA, each edge connects a TF to a target gene, and the edge weight indicates the strength of the inferred regulatory relationship.

We generated one PANDA network for each group: LCL, blood, fibroblasts, and skin (Figure S1) (Glass et al., 2013). For each network, we used the same TF/target gene prior regulatory network and the same protein-protein interaction (PPI) prior network (see below).

To generate the TF/target gene regulatory prior, we downloaded all position weight matrices (PWM) for direct and inferred *Homo sapiens* motifs from the Catalog of Inferred Sequence Binding Preferences (CIS-BP) (2015-07-07) (Weirauch et al., 2014). For each TF, we selected the motif with the highest information content, total of 695 motifs. We mapped the PWMs for these 695 motifs to promoter regions of Ensembl gene (ENSG) ids using FIMO (Grant et al., 2011). Motif mappings were parsed to only retain those below p-value cut-off of 10^−5^ and ranging from −750bp to +250bp of the transcription start site (TSS). We then converted TF motif identifiers to HGNC symbols and filtered out TFs for which no official gene symbol was known. This resulted in a regulatory prior that included 652 TFs targeting 27,249 unique Ensembl gene ids. Finally, we kept only genes that were found expressed in the GTEx filtered and normalized data set, which included 27,175 target genes.

To generate the PPI prior, we downloaded *Homo sapiens* PPI interactions (9606.protein.links.v10.txt.gz) and protein aliases (9606.protein.aliases.v10.txt.gz) from StringDb v10 (2015-10-27). We parsed this PPI data set for the 652 TFs in our TF/target gene prior. 628/652 TFs had interactions with other TFs (scores between 0-1). We added the remaining 24 TFs to our PPI prior with self-interactions only, and set all self-interactions in this PPI prior to edge scores of 1.

To run PANDA, for each sample group, we used the TF/target gene prior, the PPI prior, and the sample group gene expression data. The TF/target gene edge weights emerging from PANDA were then used to compare networks between each cell line and its tissue of origin. For pairs of networks, we compared the TF out-degree, defined as the sum of edge weights from that TF, and the gene in-degree, defined as the sum of all incoming edge weights a gene received from all expressed TFs in the network. The illustrations of the subnetworks were done using Cytoscape default yFiles Organic layout (version 3.4.0) (Shannon et al., 2003) where each edge connects a TF to a target gene, and the edge weight is represented by the thickness and color shade.

### ENCODE Chip-Seq data

Chip-Seq on GM12878 (type of LCL) targeting the TFs SMAD5, IKZF1, USF1, and USF2 were downloaded from the ENCODE Project (https://www.encodeproject.org, accessed 2016-06-03). We used the narrow peak data processed by ENCODE from 2 biological replicates (accession: ENCFF553HHF, ENCFF001VEJ, ENCFF002CIB, ENCFF001VFQ). Then, to identify genes bound by each of these TFs we used *bedtools* (v2.17) to annotate peaks that fall within the promoter region of a gene (same promoter regions used for the network reconstruction, ranging from −750bp to +250bp of the TSS).

